# An *ex vivo* lung model to study bronchioles infected with *Pseudomonas aeruginosa* biofilms

**DOI:** 10.1101/063222

**Authors:** Freya Harrison, Stephen P. Diggle

## Abstract

A key aim in microbiology is to determine the genetic and phenotypic bases of bacterial virulence, persistence and antimicrobial resistance in chronic biofilm infections. This requires tractable, high-throughput models that reflect the physical and chemical environment encountered in specific infection contexts. Such models will increase the predictive power of microbiological experiments and provide platforms for enhanced testing of novel antibacterial or antivirulence therapies. We present an optimised *ex vivo* model of cystic fibrosis lung infection: *ex vivo* culture of pig bronchiolar tissue in artificial cystic fibrosis mucus. We focus on the formation of biofilms by *Pseudomonas aeruginosa*. We show highly repeatable and specific formation of biofilms that resemble clinical biofilms by a commonly-studied lab strain and ten cystic fibrosis isolates of this key opportunistic pathogen.

## Introduction

Chronic lung infections are debilitating, highly antibiotic resistant and often lethal. They affect people with chronic obstructive pulmonary disease, ventilator-associated pneumonia, HIV/AIDS and the genetic disorder cystic fibrosis (CF). Chronic lung infections are caused by communities of different microbial genotypes and species (Short et al., 2014), but the formation of bacterial biofilms in the airways is a key factor in producing a persistent and difficult to treat infection. CF lung infections are perhaps the epitome of intractable biofilm infection:they last for decades and the majority of people with CF die from respiratory failure, 50% of them before reaching middle age (Elborn, 2016). Understanding the basic *in vivo* microbiology of key CF pathogens is a vital step towards designing effective treatment. Much research and clinical effort focuses on the opportunistic bacterial pathogen *Pseudomonas aeruginosa*, which eventually colonises most people with CF, is a primary target for antimicrobial treatment, and forms extensive biofilm plugs in the patient’s bronchioles (Bjarnsholt et al., 2009; Elborn, 2016).

Various *in vitro* systems and insect or rodent hosts are used to study *P. aeruginosa* and to determine the genetic and phenotypic variables that determine virulence and persistence. However, most *in vitro* experiments use unstructured broth cultures, or grow biofilms that are attached to abiotic surfaces and whose structure is very different from those seen *in vivo* (Roberts et al.; Bjarnsholt et al., 2013). Insect hosts have limited similarity to humans and it is now clear that rodent tissue chemistry (Benahmed et al., 2014) and immune responses (Seok et al., 2013) differ significantly from those of humans. *In vivo* experiments are also limited in duration (acute or semichronic) due to restrictions imposed by ethical concerns, cost and host response to disease. In general, the environment during chronic infection differs from that encountered in a healthy host (acute infection). Tissue damage and disease-specific changes in host phenotype cause physical differences, e.g. increased mucus volume and adhesivity in CF. The chemical environment also differs as different substrates for growth become available:in CF, bacteria use amino acids released by damaged tissues, or from mucus, as carbon sources, and do not experience the iron restriction characteristic of healthy tissue (Tyrrell and Callaghan, 2016). Consequently, gene expression and the roles played by virulence factors differ in chronic *vs*. acute contexts (Palmer et al., 2005; LaFayette et al., 2015; Turner et al., 2015). Environmental cues also affect antibiotic resistance phenotypes:*P. aeruginosa* grown in synthetic CF sputum upregulates an antibiotic efflux pump (Tata et al., 2016). A general lack of chronicity and realistic tissue chemistry therefore limits the use of *in vitro* and rodent models for investigating pathogen biology, and arguably explains the high failure rate in translating proposed new drugs from animal models to the clinic (McGonigle and Ruggeri, 2014).

Cheap, high-throughput models of lung biofilm that carefully recapitulate the physical and chemical environment encountered in chronic CF infection are conspicuous by their absence from the microbiologist’s toolkit. Such models would drastically increase the level of biological realism achievable in the laboratory and so open a new window to help us study the *in vivo* biofilm. They could be used to reveal novel targets for clinical intervention, to test promising new anti-biofilm or anti-bacterial compounds, or for more predictive diagnostic tests of antibiotic resistance.

Inspired by previously published work (Williams and Gallagher, 1978; Nunes et al., 2010), we recently developed a cheap, high-throughput protocol (Harrison et al., 2014) for infecting *ex vivo* pig lung tissue (EVPL) with *P. aeruginosa* and culturing these model infections in conditions that mimic the chemistry of chronically-infected CF lung mucus (artificial sputum medium (Palmer et al., 2007)). Pigs have more similar lung structure, immunology and chemistry to humans than do mice (Meurens et al., 2012; Benahmed et al., 2014) and lung tissue is available as a by-product from the meat industry, so the model poses no ethical concerns. The model is cheap and allows for high levels of replication:several dozen individual pieces of tissue can be dissected from each pair of lungs. Because the lungs come from animals certified fit for human consumption, the model also poses no obvious biological safety risks.

We showed that EVPL could be used to compare the growth, pathology and virulence of different genotypes of *P. aeruginosa* using cell counts, microscopy and quantitative chemical or reporter-based assays for various virulence factors (quorum sensing signals, proteases, siderophores, pyocyanin). Repeatability between tissue taken from independent lungs was high (Harrison et al., 2014). We found that, while communication via quorum sensing is required for *P. aeruginosa* growth and virulence in acute infection models, this behaviour appears to be dispensable in EVPL (Harrison et al., 2014). This shows the importance of recognising and modelling environmental and ecological differences in acute *vs*. chronic contexts.

Our previous work used sections of alveolar lung tissue, corresponding to the location infection in a very late, pre-terminal stage of CF. In reality, long-term, prophylactic use of antibiotics results in *P. aeruginosa* biofilm remaining restricted to the bronchioles for most of the course of chronic infection (Bjarnsholt et al., 2009). We have therefore developed a version of our model that uses small sections of pig bronchiole to better represent *P. aeruginosa* biofilm during the long periods of relatively quiescent chronic infection that characterise CF.

## Methods

### Bacterial strains

PAO1 and PA14 were used as examples of standard laboratory strains of *P. aeruginosa.* As exemplar chronic CF isolates, we selected ten clones taken from a single CF sputum sample that had previously been subjected to extensive phenotypic and genomic analysis in our laboratory (Darch et al., 2015).

### Artificial sputum medium and culture conditions

Artificial sputum medium was prepared following the recipe of Palmer et al. (2007), with the modification that we did not add glucose to the medium. Preliminary work suggested that glucose facilitated growth of any resident bacteria left on the lung tissue, and did not affect the growth of *P. aeruginosa* when lung tissue was present. All media used in this work were supplemented with 50 μg/ml ampicillin to further minimize the growth of any resident bacteria present in the lung tissue.

### Lung dissection & infection

Pig lungs were obtained from a local butcher (JT Beedham & Sons, Nottingham). Lungs came from the butcher’s own herd of Duroc x Pietrain pigs, were collected as soon as possible after they arrived in the shop from the abattoir and were used immediately on arrival in the laboratory. Lungs were transported in a chilled coolbox to the University of Nottingham and dissected in a room not used for microbiological work. Standard sterile technique was observed at all times (work conducted under a Bunsen burner, dissection tools pre-sterilised by autoclaving and re-sterilised as necessary by dipping in ethanol and flaming). The ventral surface of the pleura was briefly (<1 s) seared with a hot pallet knife to kill surface contaminants from the abattoir or butcher’s shop. This also renders the pleura easier to cut. A mounted razor blade was used to cut into the tissue along the length of the first 5-10 cm of the right or left main bronchus, just deep enough to expose the cartilage of the bronchus. The exact length available varies between lungs depending on the size and how rapidly the bronchus branches; we recommend working with bronchi and bronchioles of 1–3 cm in diameter. A mounted razor blade was then used to make a transverse cut across the top of the bronchus to separate it from the trachea. By holding this free end, it is possible to use the razor blade to gently separate a length of bronchus from the surrounding alveolar tissue. Approx. 5 cm long sections of bronchus/bronchiole were removed in this way, and stripped of most remaining attached alveolar tissue using mounted razor blades. These sections were washed twice with a 1:1 mix of RPMI 1640 and Dulbecco’s modified Eagle medium (DMEM) (Sigma-Aldrich). Dissection scissors were used to trim away any remaining alveolar or connective tissue on the surface of the bronchioles (this is softened and made easier to remove by washing). Scissors were then used to cut the washed bronchioles into longitudinal strips of approx. 5 mm width, and then to cut approx. 5 mm square sections from these strips. Any remaining excess alveolar or connective tissue was trimmed away with scissors as the bronchioles were being sectioned. Bronchiolar tissue sections were then washed once with a 1:1 mix of RPMI 1640 and DMEM, and once with ASM. Sections were transferred individual to the wells of 24–well tissue culture plate:each well contained a soft pad of 400 μl ASM supplemented with 0.8% agarose. 500 μl liquid ASM was added to each well. For imaging experiments only, smaller bronchioles (5 mm – 1 cm diameter) were also dissected and whole cross-sections approx. 5 mm long were cut with a mounted razor blade.

To inoculate bronchiolar sections with bacteria, a sterile hypodermic needle (29 or 30G) was lightly touched to the surface of a *P. aeruginosa* colony grown on an LB agar plate and then used to prick the bronchiolar tissue. For mock-infection controls, tissue was pricked with a sterile needle. We found that needles mounted on 1 ml insulin syringes were easy to handle safely and accurately. Tissue was incubated at 37°C on a rocking platform for up to 4 days.

After incubation, tissue was rinsed in 1 ml phosphate buffered saline to remove loosely adhering cells. Tissue sections intended for microscopy were preserved in formalin, sectioned and stained with Gram stain or haematoxylin and eosin (H&E). Microscopy was conducted with a Nikon Eclipse 50i with Digital Sight DS-U3 camera. Tissue sections used to assay total bacterial numbers were homogenised individually in 500 μl phosphate-buffered saline in metal bead tubes (Cambio) using a Precellys24 homogenizer. Homogenates were serially diluted and aliquots plated on LB agar to obtain single colonies.

### Bead biofilm assay

For each bacterial clone to be investigated, a sweep of colonies was taken from an LB agar plate, inoculated into 3ml ASM and cultured overnight at 37°C on an orbital shaker. Cultures were diluted to an OD_600_ of 0.1 in ASM and three replica 2ml aliquots transferred to 5 ml plastic universal tubes. A 9x6 mm plastic bead (pony beads from www.mailorder-beads.co.uk) was added to each tube and cultures incubated for 24 hours at 37°C on an orbital shaker at 200 rpm. Biofilms were collected by retrieving the beads from the tubes, gently washing three times in 10 ml phosphate buffered saline, transferring to 10 ml fresh phosphate buffered saline and sonicating in a bath sonicator for 10 minutes. Recovered biofilm populations were diluted and plated on LB agar to count colonies.

## Results & Discussion

Mock-infected bronchiolar EVPL retains normal histopathology for seven days (Fig. 1a). We used a sterile hypodermic needle to transfer colony-grown cells of a standard used lab strain of *P. aeruginosa*, PAO1, to EVPL and observed that after four days’ incubation in artificial sputum medium at 37°C, this strain formed dense, mucoid biofilms that are highly reminiscent of the sticky plugs that occlude CF patients’ bronchioles (Fig. 1b, c). Microscopy (Fig. 1b) showed that the biofilm had numerous empty voids, giving it a spongy appearance:this is similar to images of *P. aeruginosa* biofilm in some samples of expectorated CF sputum (Fig. 1d in Bjarnsholt et al., 2009) and in explanted CF lungs (Fig. 3 in Kragh et al., 2014), and of mucus plugs in the bronchioles of late-stage CF patients (Fig. 8 in Henderson et al., 2014). We noted that the bronchiolar tissue largely retained its integrity even when covered in large amounts of biofilm: the tissue was not dissolved or macerated by *P. aeruginosa* exoproducts.

**Figure 1a.**
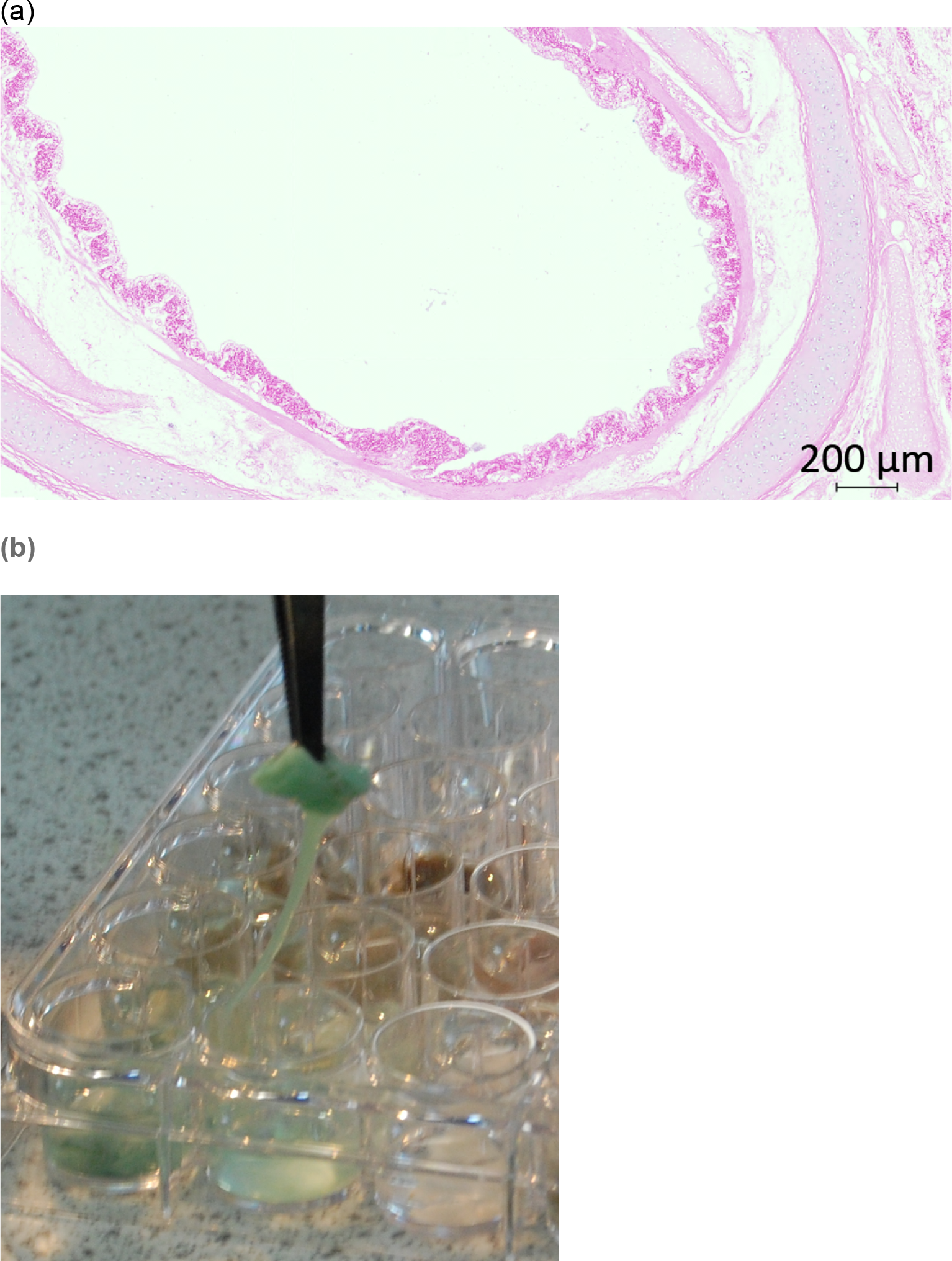
(a) Uninfected bronchiolar EVPL retains structure after 7 days’ culture at 37°C in ASM (H&E stained cross-section of a small bronchiole tissue).(b) *P. aeruginosa* strain PAO1 forms extensive mucoid biofilm on squares of bronchiolar tissue. The green pigmentation is typical of *P. aeruginosa* and is a mixture of the exoproducts pyoverdine and pyocyanin; note how the coating of bacteria drips from the tissue and sticks to the plastic culture plate as the section of bronchiole is lifted out of the well. (c) Microscopy confirms that the biofilms in (b) are a mass of Gram-negative (pink) rods.

**Figure 1b.**
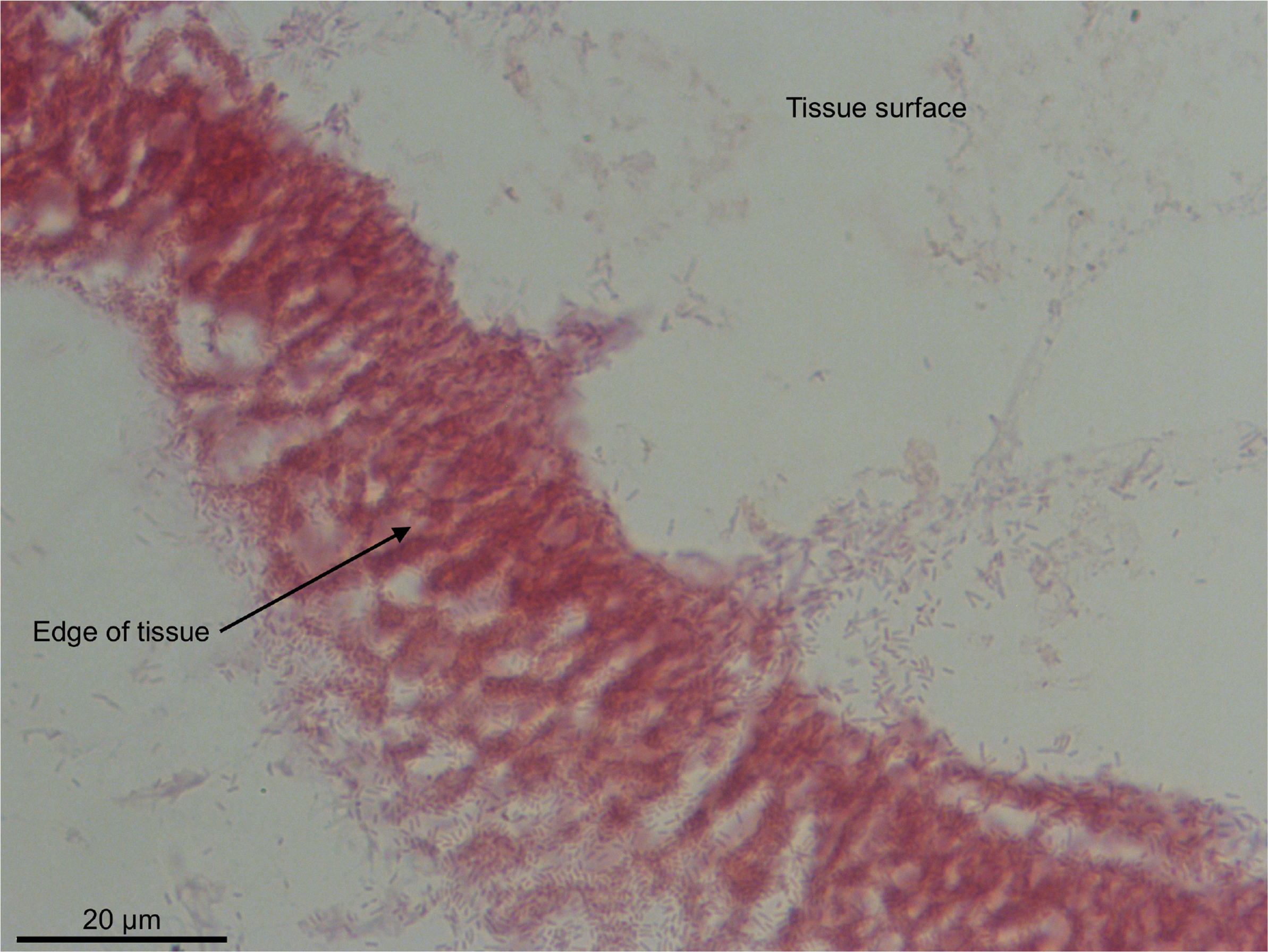

We then tested how clinical isolates of *P. aeruginosa*, that have adapted to a chronic lifestyle in CF lungs over many generations, performed in our model. We selected ten genetically and phenotypically diverse *P. aeruginosa* clones that were previously isolated in our lab from a single CF sputum sample (Darch et al., 2015) and cultured them in EVPL as described above. In parallel, we also cultured PAO1 and a second commonly-used lab strain, PA14. Three replica sections of tissue were inoculated for each strain (or used for the uninoculated control) and the whole experiment was replicated twice using two different lungs obtained on different days. As can be seen in the example photographs from lung A in Fig. 2, the CF isolates formed biofilm on bronchiolar tissue more rapidly and more specifically than the lab isolates, consistent with EVPL providing a more permissive and therefore realistic environment for these lung-adapted clones. Fig. S1 shows replica bronchiolar biofilms of each of the ten CF isolates and two lab strains at 4 days post-inoculation in lung B; comparable results were obtained in both lungs used. As with PAO1, the square of tissue retained its general shape and size, it was not destroyed by the colonising bacteria.

**Figure 2a.**
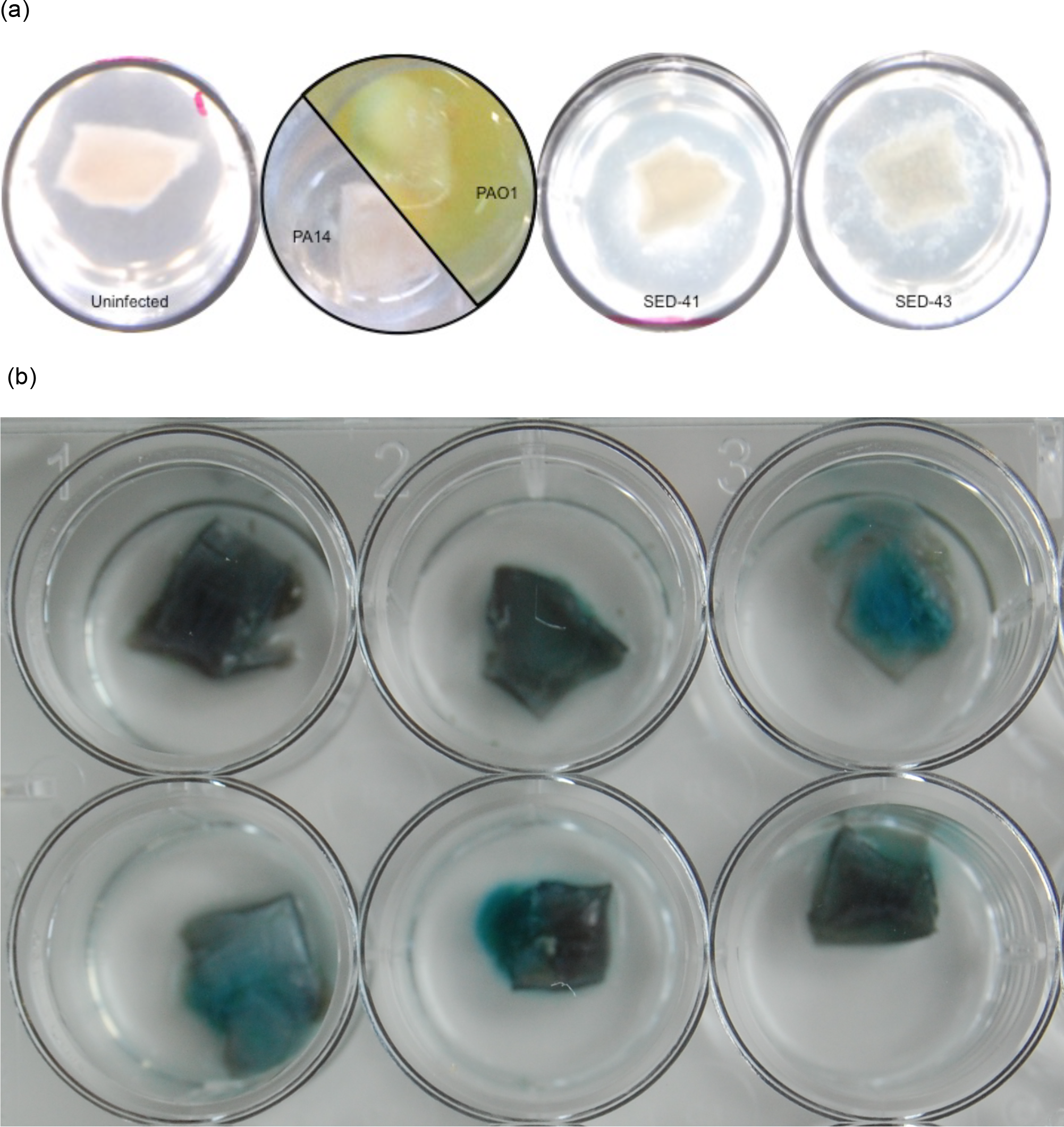
(a)EVPL *in situ* in ASM at 19 hours post-inoculation. Uninfected bronchiolar tissue retains its normal appearance:a pinkish-white square with no noticeable degradation, surrounded by clear ASM. The lab strain PA14 does not show visible growth either on the tissue or in the surrounding ASM at this early stage; PAO1, in contrast, has grown extensively in the liquid ASM surrounding the tissue (green-yellow pigmentation due to production of pyoverdine) but does not yet show any noticeable growth on the tissue itself-note pinkish-white square of tissue sitting in the liquid bacterial culture. In contrast, CF isolates of *P. aeruginosa* (e.g. SED-41, SED-43) show growth as frond-like aggregates on and connected to the cubes of tissue, very different from the dense planktonic growth of PAO1. (b) By 4 days post-inoculation, CF isolates of *P. aeruginosa* have grown to high density on EVPL. The image shows three replica infections of SED-41 (top row) and SED-43 (bottom row) after washing the tissue with phosphate-buffered saline to remove non-adhering cells:a coating of sticky *P. aeruginosa*, with blue-green pigmentation (pyoverdine and pyocyanin), is left behind. (c) These biofilms are noticeably mucoid (e.g. SED-41).

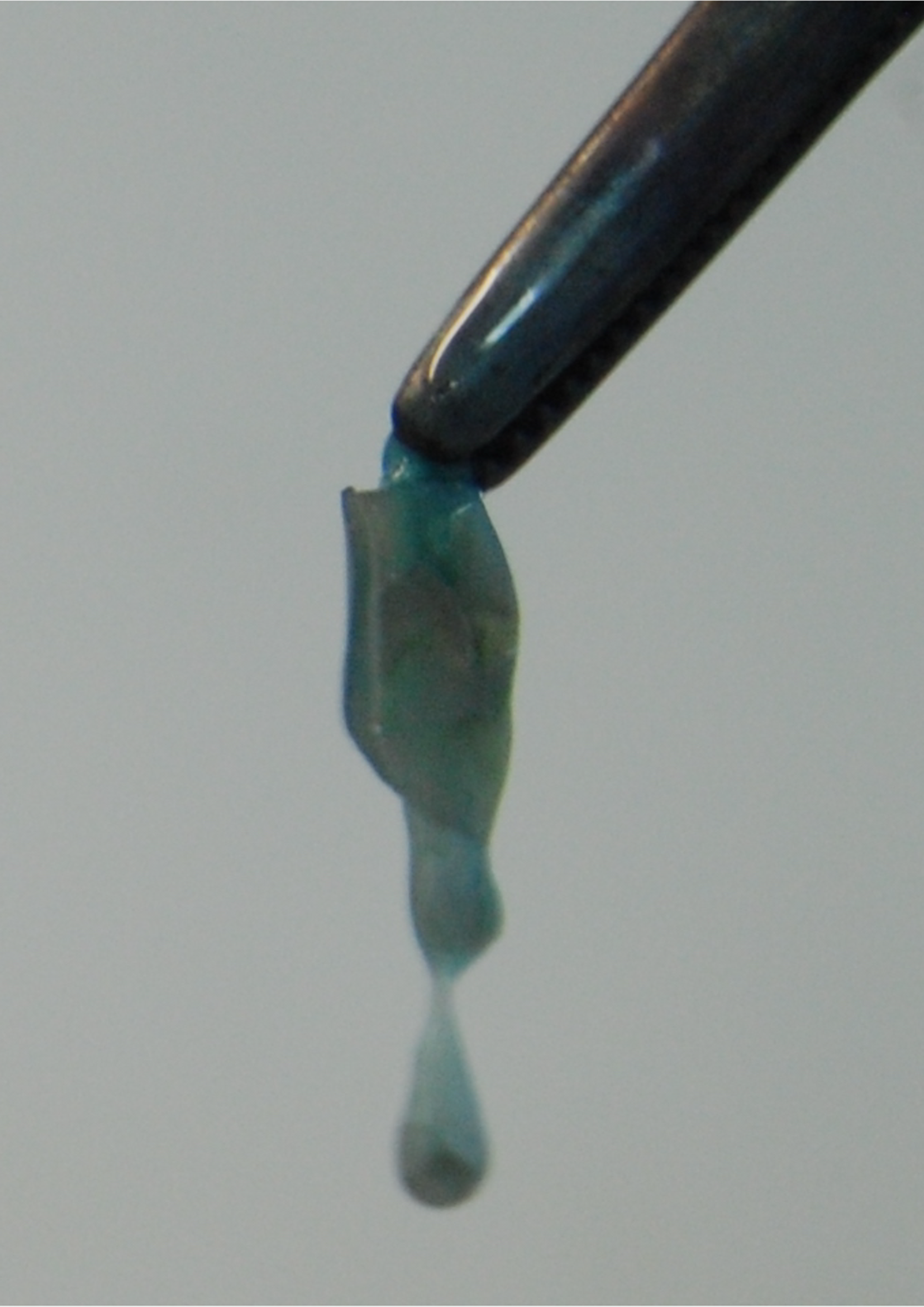

**Figure S1.**
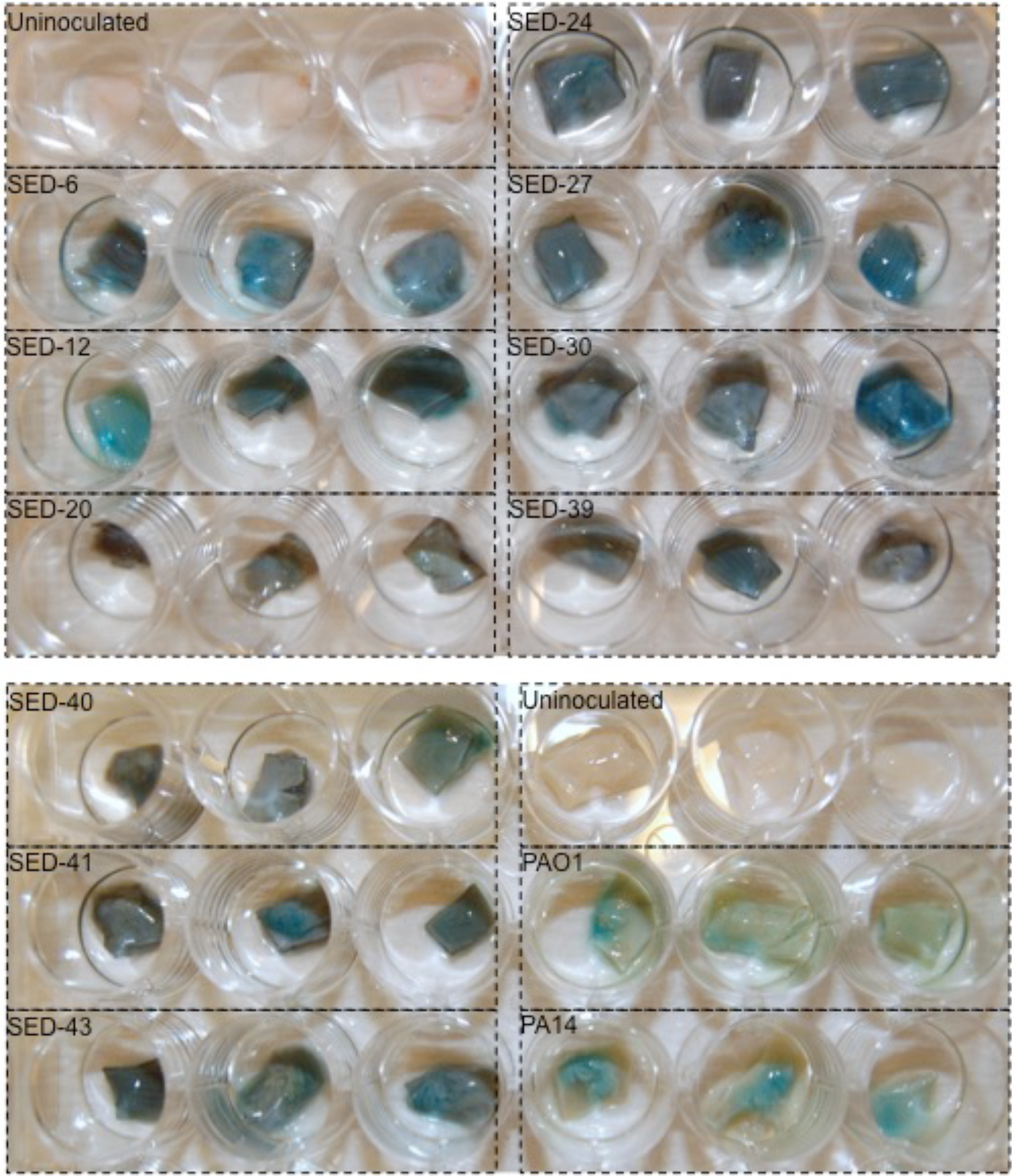
Replica biofilms formed by ten CF sputum isolates and two lab strains, shown at 4 days post-inoculation after washing in saline to remove loosely-adhering cells. Tissue is pictured in a standard 24-well culture plate; as two plates were required to grow three replicates of each clone, three replica uninoculated sections were placed in each plate. Nomenclature for the CF isolates is the same as in the first published article describing the phenotypic and genetic diversity of these isolates (Darch et al., 2015)).

It is important to determine how repeatable experimental results are likely to be in any experimental model: more variable models will require greater sample sizes to measure microbial traits of interest, or to perform experiments with adequate statistical power to reject the null hypothesis. For example, if we want to test the null hypothesis that there is no difference in biofilm forming ability between different genotypes, it would be helpful to know the proportion of variation in biofilm formation that is due to differences among clones, as opposed to differences between replica populations of the same clone. In statistical language this is called the intraclass correlation coefficient, and is easily calculated from the results of a one-way analysis of variance (ANOVA) (Lessells and Boag, 1987). Therefore, we conducted an experiment with a deliberately small sample size to compare the repeatability of biofilm formation by ten clinical isolates of *P. aeruginosa* on bronchiolar EVPL and in an attachment assay using plastic beads that has become a standard *in vitro* assay for biofilm formation (Poltak and Cooper, 2011).

Each of the ten CF isolates previously used was cultured in triplicate in EVPL (tissue from a single lung) and in a well-establised bead biofilm assay. As can be seen from the error bars in Fig. 3, there was more within-clone variation in the bead assay than in the EVPL model. Consequently, the bacterial density recovered from EVPL showed higher repeatability (0.63 vs. 0.24). This allowed ANOVA to identify inter-clone differences in biofilm formation (F_2,20_=6.1, *p*<0.001) despite the small sample size; these were not apparent in the bead assay (F_2,20_=2.0, *p*=0.104) (Table 1). The comparative data in Fig. 3 also demonstrate the impressive thickness of the biofilm formed on a relatively small surface area of bronchiolar EVPL (approx. 50 mm^2^, compared with approx. 180 mm^2^ for a bead).

**Figure 3.**
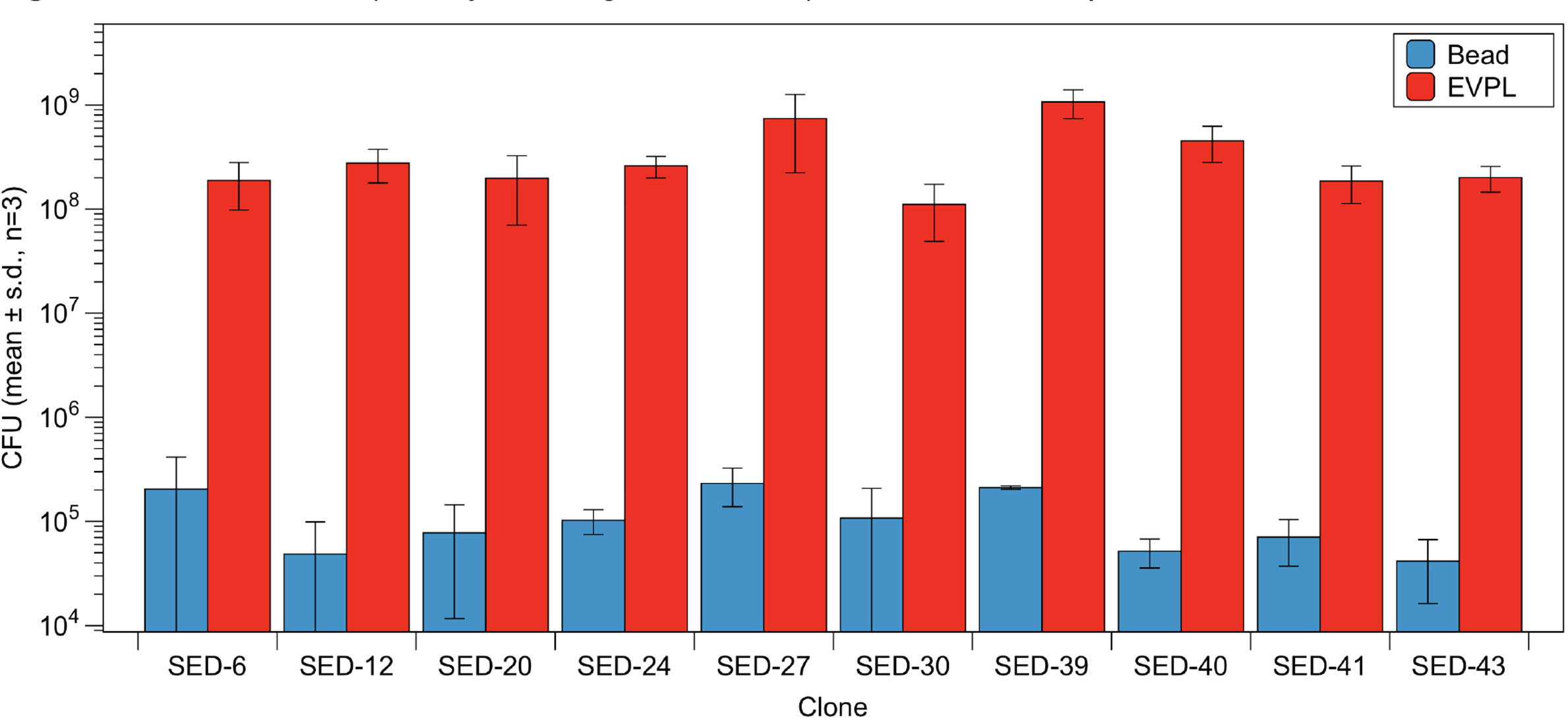
Biofilm mass (colony forming units, CFU) recovered from plastic beads and from EVPL.

**Table 1.** Analysis of Variance and repeatability (*r*) of biofilm mass for ten clinical *P. aeruginosa* isolates on (a) beads and (b) bronchiolar EVPL. Note that data for one replicate population of one clone in the bead assay was lost.

(a) Beads
| | d.f. | Sum of Squares | Mean Square | F | $p$ | $r$ |
| --- | --- | --- | --- | --- | --- | --- |
| Clone | 9 | 1.35E+11 | 1.50E+10 | 1.957 | 0.104 | 0.24 |
| Residuals | 19 | 1.46E+11 | 7.66E+09 |  |  |  |

(b) EVPL
| | d.f. | Sum of Squares | Mean Square | F | $p$ | $r$ |
| --- | --- | --- | --- | --- | --- | --- |
| Clone | 9 | 2.53E+18 | 2.81E+17 | 6.131 | < 0.001 | 0.63 |
| Residuals | 20 | 9.16E+17 | 4.58E+16 |  |  |  |

In conclusion, we present a model for CF biofilm infection that facilitates cheap, high-throughput screening of *P. aeruginosa* clones in an environment which more closely mimics the structure and chemistry of chronically-infected lungs. In our previous work with alveolar sections of EVPL, we showed that a range of bacterial virulence factors can be quantified directly from infected tissue using luminescent reporter constructs and a range of standard fluorescence-based or colorimetric assays. All of these assays will also be possible with *in situ* populations grown on bronchiolar sections, or homogenates thereof. In the future, this model will be a valuable tool in increasing our understanding of the basic microbiology of biofilm infection and its clinical consequences. We have optimised this model for the study of CF, but with a few modifications such as context-specific culture media, the model could be transferrable to the study of a range of lung infection contexts.

## Acknowledgements

We thank Johnny Pusztai for lungs; Amy Coats and Sheyda Azimi for lab assistance; Jeni Luckett, Rebecca Gabrilska and Daniella Spencer for helpful discussion of the manuscript; and the histopathology service at the Queen’s Medical Centre, Nottingham University Hospitals, for tissue preparation and staining.

